# The neuroprotective effects of Sonic hedgehog pathway agonist SAG in a rat model of neonatal stroke

**DOI:** 10.1101/2021.01.19.427193

**Authors:** Vien Nguyen, Manideep Chavali, Amara Larpthaveesarp, Srikirti Kodali, Ginez Gonzalez, Robin J.M. Franklin, David H. Rowitch, Fernando Gonzalez

## Abstract

**Objective:** Neonatal stroke affects 1 in 2800 live births and is a major cause of neurological injury. The Sonic Hedgehog (Shh) signaling pathway is critical for central nervous system (CNS) development and has neuroprotective and reparative effects in different CNS injury models. Previous studies have demonstrated beneficial effects of small molecule Shh-Smoothened-agonist (SAG) against neonatal cerebellar injury and it improves Down syndrome-related brain structural deficits in mice. Here, we investigated SAG neuroprotection in rat models of neonatal ischemia-reperfusion (stroke) and adult focal white matter injury.

**Methods:** We used transient middle cerebral artery occlusion at P10 and ethidium bromide injection in adult rats to induce damage. Following surgery and SAG or vehicle treatment we analyzed tissue loss, cell proliferation and fate, and behavioral outcome.

**Results:** We report that a single dose of SAG administered following neonatal stroke preserved brain volume, reduced inflammation, enhanced oligodendrocyte progenitor cell (OPC) and EC proliferation, and resulted in long-term cognitive improvement. Single-dose SAG also promoted proliferation of OPCs following focal demyelination in the adult rat.

**Conclusion:** These findings indicate benefit of one-time SAG treatment post-insult in reducing brain injury and improving behavioral outcome after experimental neonatal stroke.

## INTRODUCTION

Neonatal stroke (NS) results from large cerebral vessel disease and is an important cause of morbidity comprising long-term motor and/or cognitive deficits(1). With an incidence of 1 in 2800 live births, NS is more common than adult large vessel stroke. Secondary injury following arterial ischemia and reperfusion can lead to cell death, oxidative stress, inflammation, and altered cell proliferation and fate(2). Cerebral white matter is also highly vulnerable to ischemic injury during the neonatal period, resulting in significant functional impairment(3), which in part reflects adverse effects on oligodendrocyte lineage cells. Indeed, oligodendrocyte progenitor cells (OPCs) continue to undergo significant changes during the perinatal and postnatal period, and are vulnerable to ischemic injury(4). Currently we lack neuroprotective therapies for NS.

Sonic hedgehog (Shh) signaling plays a crucial role in early CNS development, influencing oligodendrocyte development(5). Shh is a secreted protein that functions by inhibiting the transmembrane receptor Patched and inducing activity of Smoothened, allowing for activation of downstream pathway targets including Gli1 and Cyclin D1 (CCD1). The cilium is present on neural stem cells and has recently been shown to be present in OPCs but absent in mature oligodendrocytes, corresponding to loss of responsiveness to Shh(5). In addition, astroglial-secreted Shh has been shown to play a role in blood-brain-barrier (BBB) maturation and maintenance, along with post-insult anti-inflammatory effects(6).

SAG is a small molecule agonist of the Smo-Shh pathway. We have previously demonstrated efficacy of SAG treatment in a mouse model of neonatal cerebellar injury(7). In addition, SAG treatment starting at birth has also been found to be beneficial in a mouse model of Down syndrome(8). However, these models are distinct from the forebrain injury commonly seen in human preterm and full-term neonates.

We have previously described a non-hemorrhagic ischemia-reperfusion focal stroke model in the P10 immature rat using transient unilateral middle cerebral artery occlusion (tMCAO)(9). tMCAO mimics large vessel neonatal stroke and is thus suitable for testing pathobiology and post-injury treatment strategies. For instance, we have described alterations in neural precursor cell fate following tMCAO that affects both the subcortical white matter and cortical/subcortical gray matter(9). Oligodendrocyte injury leads to loss of myelin sheaths, which are important for metabolic support and transmission of action potentials(10). Oligodendrocyte maturation arrest is observed in neonatal white matter injury(11). Therefore, enhancing and optimizing the endogenous oligodendrocyte response could improve long-term outcomes. To further elucidate oligodendrocyte biology following demyelinating injuries, we used the adult cerebellar peduncle focal demyelination model as it has been well characterized(12).

There has been a significant focus on developing therapeutic strategies using biologics to enhance endogenous repair processes following brain injury. Recent studies have shown preserved neuronal numbers, angiogenesis and improved outcomes following Shh or SAG treatment in adult stroke models(13). Here we investigated neuroprotective effects of SAG in two distinct models of focal brain injury: (1) neonatal unilateral MCA ischemia-reperfusion in the immature rat, and (2) focal toxin-induced demyelinating cerebellar peduncle white matter injury in the mature rat. We examined both short-term histological and long-term behavioral outcomes after early stroke, and SAG-mediated cell type-specific effects on forebrain gray and white matter parenchyma.

## METHODS

All animal research was approved by the University of California-San Francisco and University Biomedical Services of the University of Cambridge Institutional Animal Care and Use Committees, and in compliance with United Kingdom Home Office regulations.

### Preparation of SAG and Vehicle

Synthesis of SAG was performed as described previously(14). SAG was dissolved to 5mM in dimethyl sulfoxide (DMSO) then further diluted in normal saline to contain 5% DMSO.

### In vivo SAG bioactivity

SAG was injected in *Gli-Luciferase* reporter mice(15) at a concentration of 50mg/kg body weight. Mice were anesthetized with isoflurane and placed in an *in vivo* bioluminescent imaging system (Xenogen) to detect luciferase reaction. SAG bioactivity was visualized by increase in intensity in transgenic animals.

### Transient middle cerebral artery occlusion

Postnatal day 10 (P10) Sprague-Dawley rats underwent focal ischemia-reperfusion with transient right middle cerebral artery occlusion (tMCAO) for three hours, or sham surgery, as previously described(9). The right internal carotid artery (ICA) was dissected and an arteriotomy was made proximal to the isolated ICA. A silicone coated 6-0 nylon filament was inserted 9-10.5 mm to occlude the MCA and secured for the duration of occlusion. Following induction, rectal temperature was monitored and maintained at 36 °C–37 °C with a combination of heating blanket and overhead light until recovery from anesthesia. For reperfusion, each animal was anesthetized, and suture ties and the occluding filament were removed. Avitene Microfibrillar Collagen Hemostat was placed over the arteriotomy and the skin incision was closed. Sham animals were anesthetized, and the ICA was dissected, after which the skin incision was closed. Immediately following reperfusion animals received a single intraperitoneal (IP) injection of SAG (50 mg/kg), or vehicle. Weight was monitored for one week following tMCAO or sham surgery to ensure adequate weight gain. 32 animals underwent tMCAO, and 22 animals sham surgery (54 animals total). Animals were split into 4 groups (Sham+Veh, n=5; Sham+SAG, n=5; tMCAO+Veh, n=8; tMCAO+SAG, n=8, Supplementary Figure S1) for cell fate analysis at two weeks post-surgery, and then were injected with BrdU (25 mg/kg twice daily) from age P11-P15 to label newly generating cells. For the second cohort of animals, behavioral analysis was performed at two months of age (Sham+Veh, n=6; Sham+SAG, n=6; tMCAO+Veh, n=8; tMCAO+SAG, n=8).

### Western blot analysis and quantitative reverse transcription-PCR

Tissue lysates were obtained from ipsilateral and contralateral cortices of Sham+Veh, Sham+SAG, tMCAO+Veh, and tMCAO+SAG 24hours post-reperfusion or equivalent (n=3 per group). For protein analysis, fresh-frozen tissues were homogenized in RIPA buffer (Thermo Scientific) containing protease inhibitors (Roche). Immunoblotting and fluorescent detection were done as previously described using the Odyssey system (Li-Cor)(4). Antibodies included Cyclin D1 (rabbit polyclonal, Thermo), Iba1 (rabbit polyclonal, Wako), GFAP (mouse monoclonal, Sigma), and β-Actin (mouse ascites, Sigma). For qRT-PCR, RNA was isolated with Trizol followed by RNeasy (Qiagen) from tissue lysates and assayed for *Gli1* gene expression with SYBR-Green (Roche) on a LightCycler 480 (Roche).

### Histology

Animal brains were harvested at either two weeks post-surgery, or following completion of behavioral testing at 2 months, as previously described(16). The entire brain was sectioned at 50-μm intervals on a sliding microtome (Thermo), and serial sections were mounted, dried, and stained with Cresyl Violet or the following antibodies: BrdU [Rat Mab BU1/75(ICR1), Accurate Chemicals], Olig2 (Mouse Mab from C.D. Stiles, Harvard), GFAP (Rat Mab 2.2B10, Thermo), Iba1 (Rabbit Pab, Wako), CC1 (APC, Mouse Mab OP80, Calbiochem), Nkx2.2 (Mouse Mab 74.5A, Developmental Studies Hybridoma Bank), Collagen 4a (Goat Pab, AB769 Millipore), Claudin 5 (Mouse Mab, 35-2500 Invitrogen).

### Focal demyelination in cerebellar white matter

Young CD rats (10-12 weeks old, Charles River) were used for the surgical procedures. Ethidium bromide (EtBr, 0.01%) was injected into the rat caudal cerebellar peduncle (CCP) bilaterally to induce demyelination as previously described(12). Animals in the treatment group received SAG (50 mg/kg) IP 5 days post injury (dpi) while control (sham) group received a single injection of saline IP at the same time point. OPC recruitment, proliferation and differentiation are key for successful remyelination. To avoid the acute effects in this model on OPC proliferation and recruitment and focus on later repair, treatment was administered 5 days following EB injection as previously published (17, 18). Treatment and sham animals were perfused at 7 or 14 dpi to study histology. Frozen sections (12μm) were collected using a cryostat (Bright Instruments). Oligodendrocyte proliferation and maturation were studied at 7 dpi (Nkx2.2, Sham+Veh, n=4, Sham+SAG, n=3; Nkx2.2/Ki67, Sham+Veh, n=4, Sham+SAG, n=3; Olig2/CC1, Sham+Veh, n=4, Sham+SAG, n=4; PLP, Sham+Veh, n=3, Sham+SAG, n=4), while mature oligodendrocytes were studied at 14 dpi (Olig2/CC1, Sham+Veh, n=5, Sham+SAG, n=5; PLP, Sham+Veh, n=4, Sham+SAG, n=3; Supplementary Figure S1).

### In situ hybridization

The expression of Plp mRNA (1:2000) in demyelinated lesions was examined by *in situ* hybridization with digoxigenin-labelled cRNA probes on cryostat sections (12 μm) using established protocols(19). ImageJ 1.44 (Wayne Rasband) was used to determine the number of positive cells within the lesions on digitized sections.

### Behavioral testing

Cylinder Rearing (CR) was used to assess the effects of ischemic injury and SAG treatment on forelimb use as a function of sensorimotor bias, as previously described(20). Animals were handled for about 5 minutes per day for three days prior to testing, which was conducted 7.5 weeks after surgery. Each animal was then individually placed in a transparent Plexiglass cylinder (20-cm diameter, 30-cm height) and observed for 3-minute trial on two consecutive days, with results averaged per animal. Initial forepaw placement of each weight-bearing contact with the wall was recorded as right, left, or both forepaws. Results were expressed as the percentage use of the non-impaired (right) forepaw for braces relative to the total number of forepaw initiations.

Novel object recognition (NOR) was performed to assess recognition memory at 8 weeks following surgery(21). Habituation was performed on days 1 and 2, for a 10-minute trial each day, in a 40 × 40cm box with 30cm tall walls in a room lacking distinctive navigational cues. A 5-minute familiarization trial was performed on day 3, with two identical objects inside the testing box. Finally, a 3-minute testing phase or NOR trial was performed on day 4, 24 hours following the familiarization phase, where one familiar object was replaced by a new object of the same size. The animal exploration time of the novel object was recorded: novel object preference index (%) = novel object exploration time / (novel object recognition time + familiar object exploration time) × 100.

### Stereological analysis

Unbiased, systematic random sampling was used to quantify both hemisphere and corpus callosum volume. A series representing every 12^th^ section was selected, Cresyl Violet stained, and analyzed. Sections encompassed the whole brain rostrally from the genu of the corpus callosum through the posterior portion of the hippocampus to the occipital lobes caudally. All volumetric quantifications were performed in a blinded manner on a Zeiss AxioScope Imager Z.2 with a motorized stage, with Neurolucida and Stereoinvestigator software (MicroBrightField). For the ROIs, the ipsilateral (lesioned) and contralateral (control) hemispheres (4x magnification) and corpus callosum (10x) were traced. The cross-sectional area of the region of interest (ROI) was calculated according to the Cavalieri principle, as previously published(22). Damage secondary to stroke was determined quantitatively by calculating the percent volume of the ipsilateral versus contralateral hemispheres.

For quantitative analysis of cell types, a systematic random sampling of every 24^th^ coronal section (3-4 sections per animal) was single or double-immunostained. Sections were imaged using a Zeiss AxioImager.A2 upright microscope and analyzed using Stereoinvestigator. On each section, a tile scan of the entire hemisphere was imaged, the ROI (striatum or corpus callosum) was outlined, and the 20X field was randomly placed within the ROI for imaging. Following imaging of the full thickness z stack (1-μm steps) with 10-μm border zones, the field was moved in an automated fashion resulting in imaging of approximately 18% of the ROI. Single and co-labeled cells were quantified using the Optical Fractionator probe for unbiased cell counting within the software. Cell density was calculated as average number of cells per 20X field, cell percentage was calculated as total number of co-labeled cells per single label, and total cell number in the corpus callosum was calculated using Nv x Vref calculation based on density and volume of the measured corpus callosum, respectively.

### Statistical analysis

Data are presented as mean ± SEM. Sex was equally distributed between groups and time points, and no differences were seen between female and male animals. All analyses were conducted by investigators blinded to treatment group. For statistical analysis, nonparametric methods were used. The one-way ANOVA Kruskal-Wallis test was applied for comparisons of multiple groups followed by the Wilcoxon rank sum test for comparison between two groups. P values were corrected by the Bonferroni method to maintain α value at 0.05. All statistical analyses were performed using SAS Enterprise Guide software (SAS Institute, Cary, NC, U.S.A.).

## RESULTS

### Single-dose SAG treatment preserves brain volume following neonatal tMCAO

We determined the optimal concentration of SAG (50 mg/kg) by dose-response *in vivo* using Gli-luciferase mice injected at P10 and followed for 120 hours (Supplementary Figure S2), where we saw elevated luciferase signal for at least 72 hrs. We tested different doses in rats (10 to 100mg/kg body weight, data not shown), and found that 50mg/kg resulted in the most potent changes similar to our results in mice. To determine the efficacy of post-insult one-time SAG treatment on forebrain injury in the immature brain, we performed 3-hour tMCAO or sham surgery in P10 rat pups who were then treated with SAG or vehicle at the time of reperfusion (Fig. 1A). In rats, P10 is considered a rough developmental equivalent to the full-term human neonate, when NS is most common. Prior work has shown that the tMCAO model consistently results in moderate injury involving the ipsilateral striatum, subcortical white matter and parieto-temporal cortex(9). Both sham-operated animals that received SAG or vehicle treatment (Sham+Veh = 0.98±0.03, Sham+SAG = 1.01±0.04; Fig. 1B, C) showed minimal hemispheric impact. While tMCAO+Veh animals showed consistent, significant hemispheric volume loss two weeks following tMCAO (tMCAO+Veh = 0.61±0.04; p<0.001) single-dose SAG treatment at time of reperfusion significantly preserved hemispheric brain volume in tMCAO animals (tMCAO+Veh = 0.61±0.04, tMCAO+SAG = 0.82±0.03; p<0.01), including the corpus callosum (CC) (tMCAO+Veh = 0.565 ± 0.04; tMCAO+SAG = 0.732 ± 0.05, p<0.05, Fig. 1D), almost to sham levels. Thus, there is a significant neuroprotective effect of a one-time SAG treatment following a tMCAO ischemic insult.

**Figure 1:**
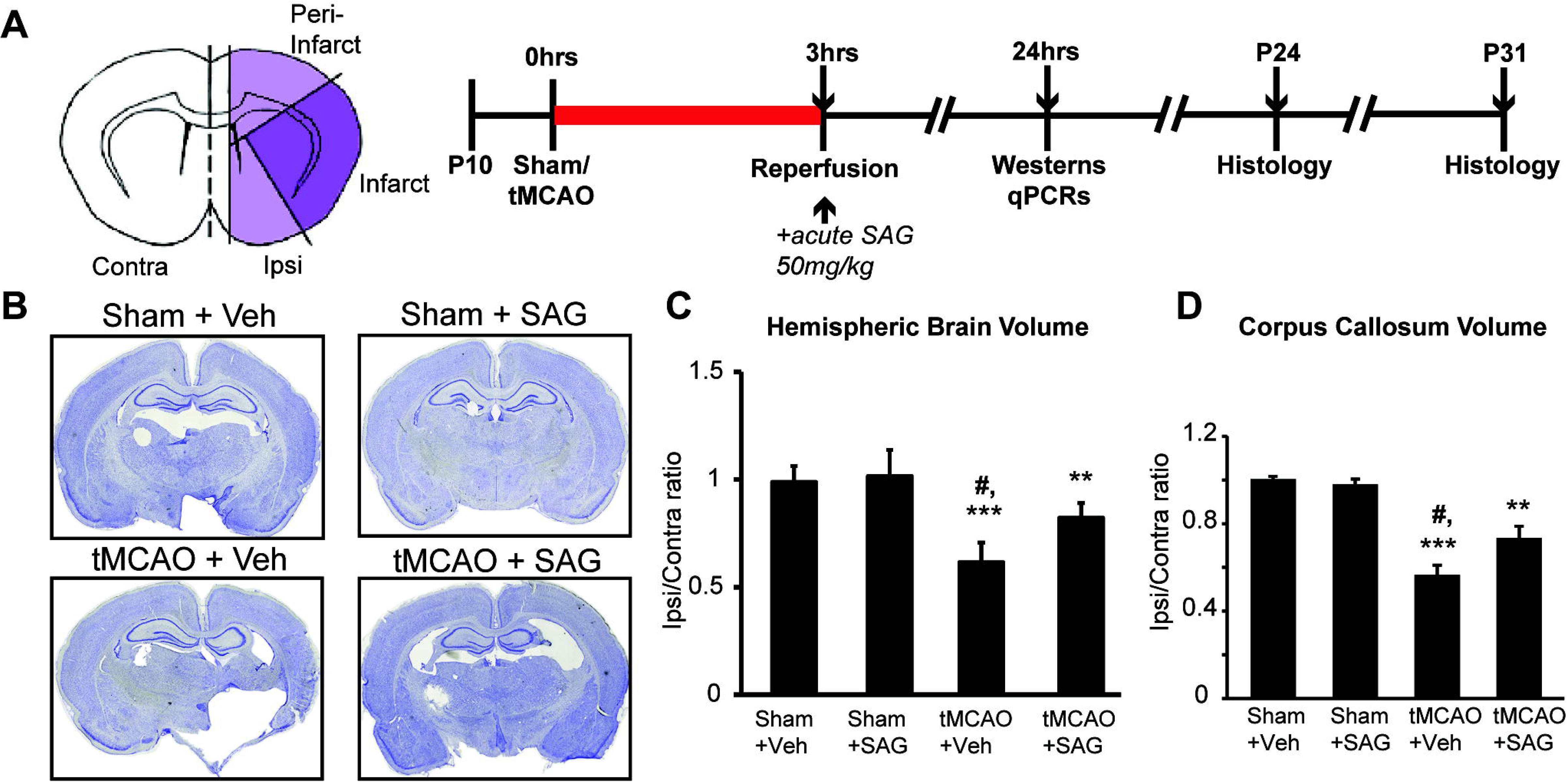
SAG administration following tMCAO improves brain volume. (A) Diagram of transient middle cerebral artery occlusion (tMCAO) injury region in a coronal section of a rat brain, and experimental timeline for tMCAO and volume/cell fate analysis. Contra, contralateral/uninjured hemisphere; Ipsi, ipsilateral/injured hemisphere. (B) Representative images of coronal sections from rat brains two weeks post-tMCAO or sham surgery, at P24. (C) Quantification of brain volume following surgery and treatment, showing ratio of injured (ipsilateral) and uninjured (contralateral) hemispheric volumes. (D) Quantification of corpus callosum volume following surgery and treatment. For quantification, mean ± SEM; **p < 0.01 and ***p < 0.001 vs Sham+Veh, #p < 0.05 vs tMCAO+SAG; one-way ANOVA with Dunnett’s test. Veh, injection of vehicle solution (0.85% saline solution).

### SAG treatment activates downstream Shh targets in the ipsilateral hemisphere and reduces inflammation following neonatal tMCAO

To confirm that SAG administration targets the Shh pathway, we performed protein/gene expression analysis from brain samples collected 24h after reperfusion to determine expression of downstream Shh targets cyclin D1 (CCD1) and Gli1. Both CCD1 (Fig. 2A, B) and Gli1 (Fig. 2C) showed significantly increased protein (CCD1) and gene expression (Gli1) levels in the tMCAO+SAG group (CCD1 Ipsi/Contra expression ratio: tMCAO+Veh= 0.60±0.047, tMCAO+SAG=1.5 ±0.26, p<0.05; Gli1 Ipsi/Contra expression ratio: tMCAO+Veh=0.24±0.0014, tMCAO+SAG=1.1±0.17, p<0.05).

**Figure 2:**
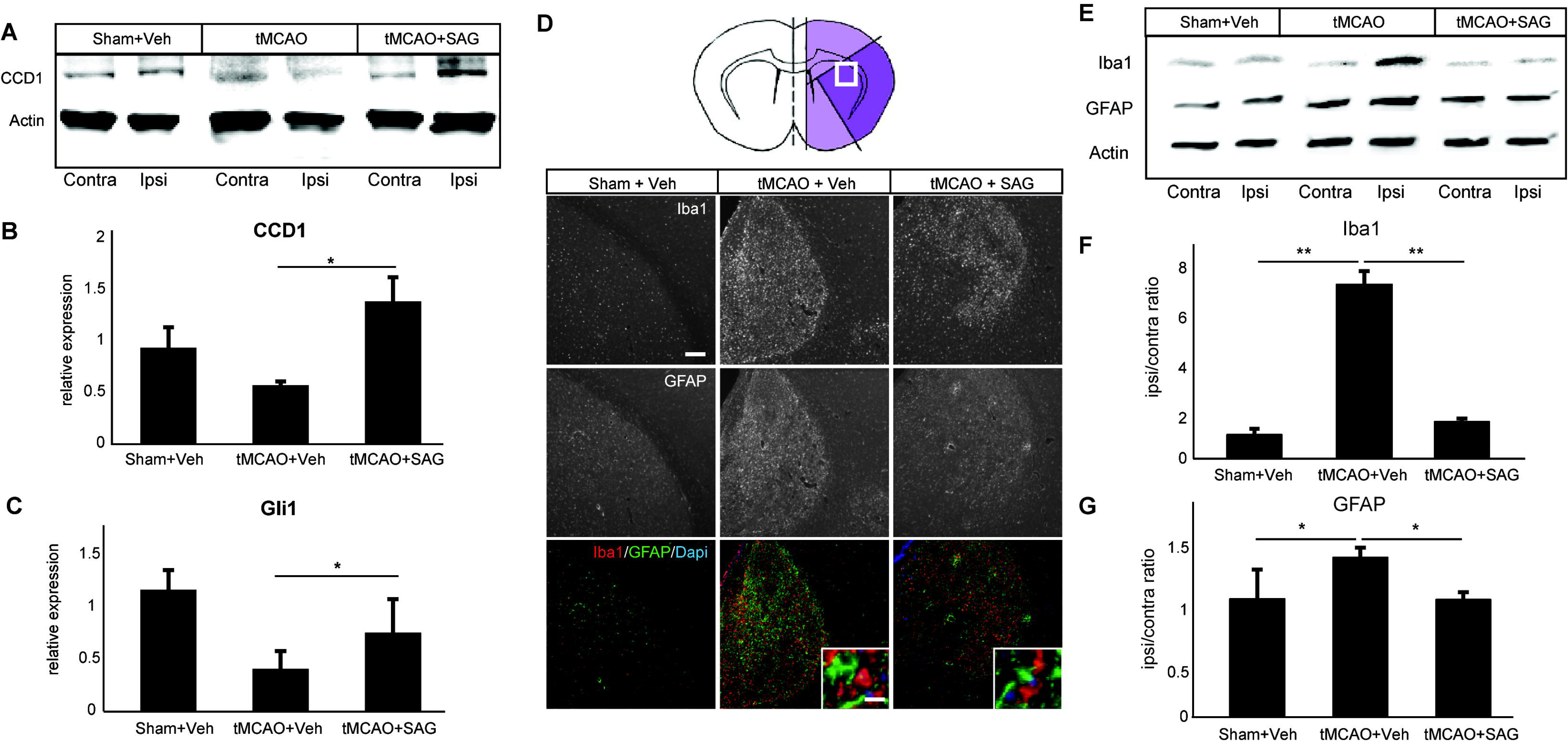
SAG activates downstream Shh targets and decreases gliosis. (A) Brains collected 24hrs post-surgery analyzed for protein expression of the Shh downstream target Cyclin D1 (CCD1) expression in Sham+Veh, tMCAO+Veh, or tMCAO+SAG. Contra, contralateral/uninjured hemisphere; Ipsi, ipsilateral/injured hemisphere. (B) Quantification of CCD1. (C) tMCAO brains collected 24hrs post-surgery analyzed for mRNA expression of Shh downstream target *Gli1* using qRT-PCR. (D) Diagram showing striatum from infarct region of tMCAO brain, and representative immunofluorescence images of Iba1, GFAP, or merged. Insert, higher magnification of cells positive for Iba1 and GFAP. Scale bar, 500μm; inset, 20μm. (E) Representative immunoblots of Iba1 and GFAP expression in Sham+Veh, tMCAO+Veh, or tMCAO+SAG brains. Contra, contralateral/uninjured hemisphere; Ipsi, ipsilateral/injured hemisphere. (F) Quantification of Iba1 ipsilateral-to-contralateral expression ratio. (G) Quantification of GFAP. *, p < 0.05; **, p<0.005, Student’s t-test.

Neonatal hypoxia-ischemia injury is typically followed by widespread inflammation involving parenchymal microglia and astrocytes, which could contribute to secondary injury(23). To investigate the effects of SAG treatment on gliosis we performed immunohistochemical analysis of activated microglia (Iba1)) and reactive astrocytes (glial acidic fibrillary protein, GFAP) in grey matter regions post-neonatal tMCAO. As shown (Fig. 2D, E), we observed a decrease in ipsilateral Iba1 staining, which suggests decreased microglial inflammation, and astrocytosis in tMCAO+SAG compared to tMCAO+Veh animals. When quantified by immunoblotting, both Iba1 and GFAP protein expression were significantly increased in the ipsilateral hemisphere in tMCAO+Veh compared to sham groups, but showed no significant difference when SAG was administered (Sham+Veh = 1.05 ± 0.26; tMCAO+Veh = 7.58 ± 0.59; tMCAO+SAG = 1.55 ± 0.13, p<0.01. Fig. 2F, G). These findings indicate that SAG treatment induces Shh targets and decreases inflammation following tMCAO.

### SAG increases OPC proliferation following neonatal tMCAO and adult demyelinating injury

Our analysis that SAG could improve hemispheric brain volume preservation specifically showed improvements in the corpus callosum (CC), which could reflect axon preservation and/or support of myelinating oligodendrocytes or OPCs. We next focused on effects of SAG treatment on the oligodendrocyte lineage. Using Olig2 as a pan-lineage marker and BrdU+ co-labeling, we observed a significant increase in proliferating OPCs in SAG-treated animals 14-days post-tMCAO in the ipsilateral corpus callosum (tMCAO+Veh = 769 ± 371 vs. tMCAO+SAG = 3400 ±1235; p<0.01, Fig. 3A, B), but not total Olig2+ cells (Fig. 3C). This was associated with an increase in the proportion of NKX2.2+ OPCs in the tMCAO+SAG group compared to controls (Shams = 0.052 ± 0.023; tMCAO+Veh = 0.16 ± 0.023; tMCAO+SAG = 0.27 ± 0.18; p<0.01, Fig. 3D, E). However, despite increased immature oligodendrocytes in the CC of SAG-treated animals, we found that myelin basic protein (MBP) labeling appeared similar to Veh-treated controls (Fig. 3F), and that co-labeled mature Olig2/CC1+ oligodendrocytes were similar in the two groups (Fig. 3G, H).

**Figure 3:**
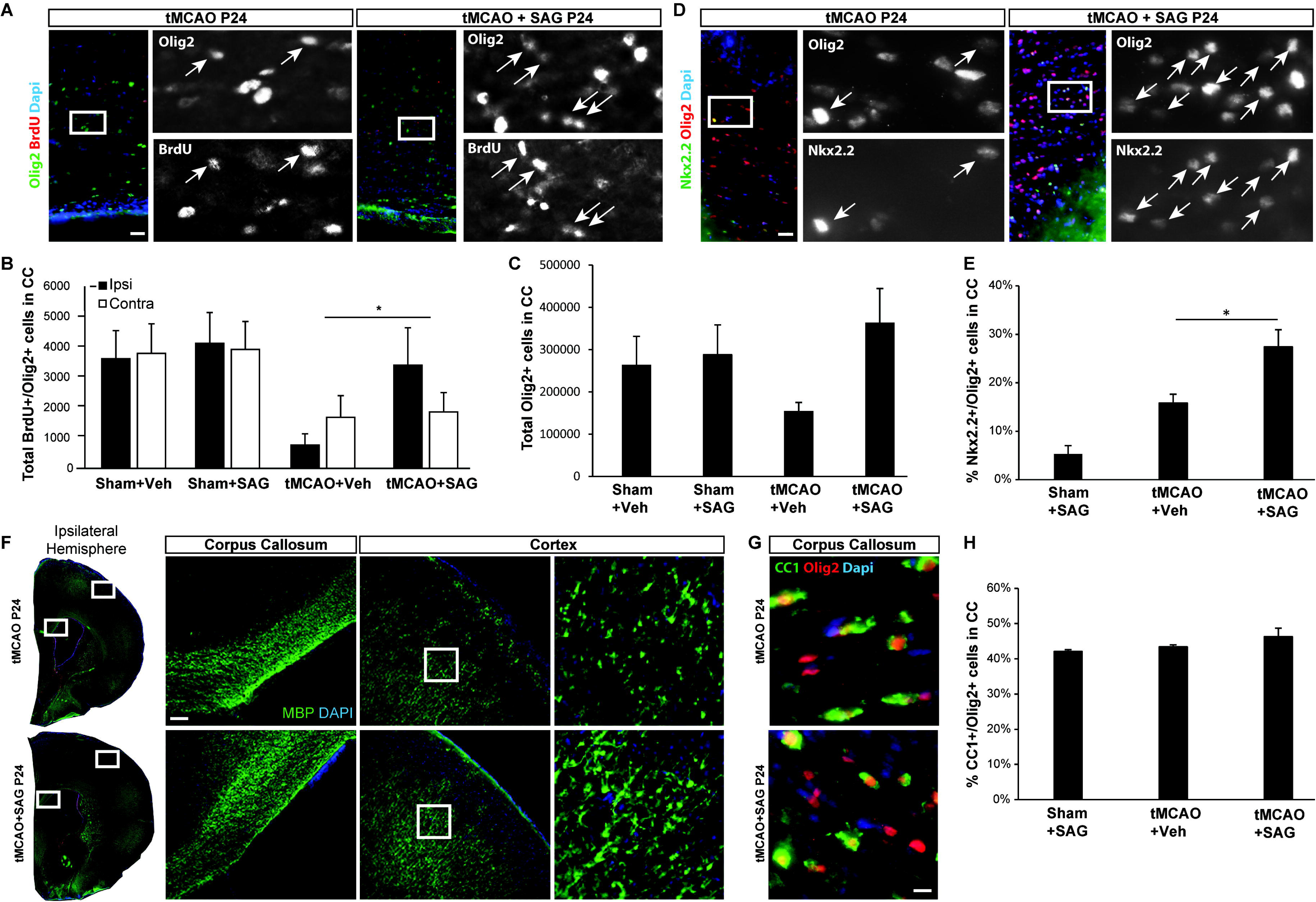
SAG increases oligodendrocyte proliferation but not differentiation following tMCAO. (A) Representative images showing proliferating cells (BrdU+) in the corpus callosum (CC) positive for Olig2 in rats given tMCAO administered with saline (left) or SAG (right) at P24. Scale bar, 50μm. Insert, magnified view of single-channel immunostains. Arrows denote double-labeled cells. (B) Quantification of cells double-labeled with BrdU and Olig2 in the CC. Black bars, CC from ipsilateral hemisphere; white bars, CC from contralateral hemisphere. (C) Quantification of total Olig2+ cells in the CC in the ipsilateral hemisphere. (D) Representative images showing Olig2+ cells double-labeled with Nkx2.2 in the CC in rats given tMCAO administered with saline (left) or SAG (right) at P24. Scale bar, 50μm. Insert, magnified view of single-channel immunostains. Arrows denote double-labeled cells. (E) Proportion of Nkx2.2+/Olig2+ cells in the CC of the ipsilateral hemisphere in tMCAO+SAG versus tMCAO+Veh. (F) Representative images of the ipsilateral hemispheres stained for MBP (green) in P24 rats (left), and inserts shown at higher magnification (right). Scale bar, 500μm. (G) Representative images showing Olig2+ cells double-labeled with CC1 in the CC in rats given tMCAO administered with saline (top) or SAG (bottom). Scale bar, 20μm. (H) Quantification of CC1+/Olig2+ cells in the CC of the ipsilateral hemisphere in tMCAO+SAG versus tMCAO+Veh. *, p < 0.05, one-way ANOVA with Tukey’s multiple comparisons test.

To further interrogate SAG effects on remyelinating oligodendrocytes, we used a focal ethidium bromide (EB) induced demyelination model(12) in adult rats followed by one-time SAG or vehicle injection (Fig. 4A, B). This model is characterized by demyelination without axonal loss and has a well-established temporal sequence of remyelination involving OPC proliferation (7 dpi) and differentiation (14 dpi). The lesion area was determined by DAPI hyper-cellularity, and we used NKX2.2/Ki67 and Olig2/CC1 to mark proliferating OPCs and differentiated cells, respectively. As shown (Fig. 4C, D), at 7 dpi there were increased numbers of newly generated OPCs (Veh = 11 Nkx2.2 cells/mm^2^; SAG = 20 cells/mm^2^, p<0.05). There was an increase in the number of Olig2/CC1 at 7 dpi which suggests a pro-differentiation effect of SAG on OPCs at early time points; there was no differences at 14 dpi between vehicle and treated group. Staining for the Proteolipid protein (Plp), a marker of myelin-forming oligodendrocyte, showed an increase of the Plp+ cells: however, this was not statistically significant (similar results were found at 14 dpi tissue; there were no statistically significant differences between groups). Together these findings indicate that SAG treatment transiently increases oligodendrocyte proliferation at early stages post-injury, but has no sustained effects on differentiation and remyelination. Taken together, these findings suggest SAG treatment results in increased CC volume and levels of OPC proliferation but not oligodendrocyte differentiation or remyelination.

**Figure 4:**
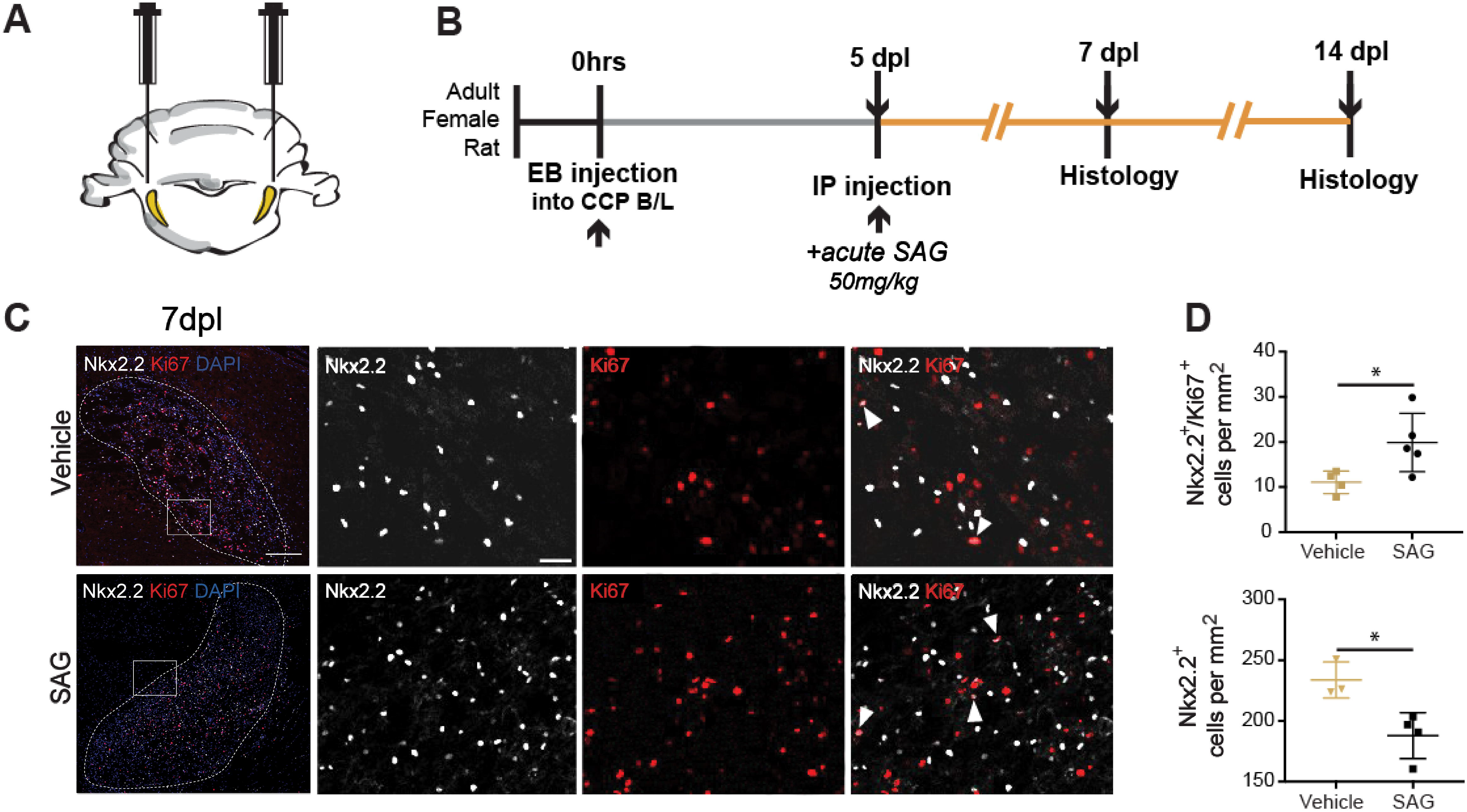
SAG promotes oligodendrogenesis and maturation in an adult rat model of focal demyelination. (A) Schematic of EB model. (B) Experimental timeline for focal demyelination using ethidium bromide (EB) injection into the rat cerebellar peduncle (CCP). (C) Representative images of proliferating cells (Ki67, red) and OPCs (Nkx2.2, white) stained within the lesioned area (dotted lines) at 7 days post injury (dpi). Scale bar, 25μm. Insert, magnified view. Scale bar, 10μm. Arrowheads denote cells double-positive for Ki67 and Nkx2.2. (D) Quantification of Nkx2.2+/Ki67+ cells (top), and total Nkx2.2+ cells (bottom) from lesioned areas after vehicle or SAG administration. * p < 0.05, Student’s t-test.

### SAG promotes angiogenesis following tMCAO

Angiogenesis is a universal feature of wound healing across tissues, and Shh has known effects in promoting vascular integrity. To determine if SAG treatment promoted vascular remodeling post-injury, we co-labeled sections harvested at P24 with BrdU+ (proliferation) and Collagen 4A (Col4A; a vascular basement membrane marker), to quantify endothelial cell (EC) proliferation in the injured core following tMCAO or sham surgery (Fig. 5A). As OPCs have been shown to play an important role in endothelial proliferation and vascular development during cerebral development(24) and under hypoxic conditions(4), we also explored the possibility that an increase in OPC number is associated with angiogenesis following injury and treatment. We did observe a close association of proliferating oligodendrocytes with the vasculature in the injured striatum in tMCAO+SAG compared to tMCAO+Veh animals (Fig. 5B). Finally, immunohistochemical analysis of Claudin-5 co-labeled with Col4A indicated no significant change in expression or vascular coverage, suggesting that SAG does not regulate endothelial tight junction protein expression (Fig. 5C).

**Figure 5:**
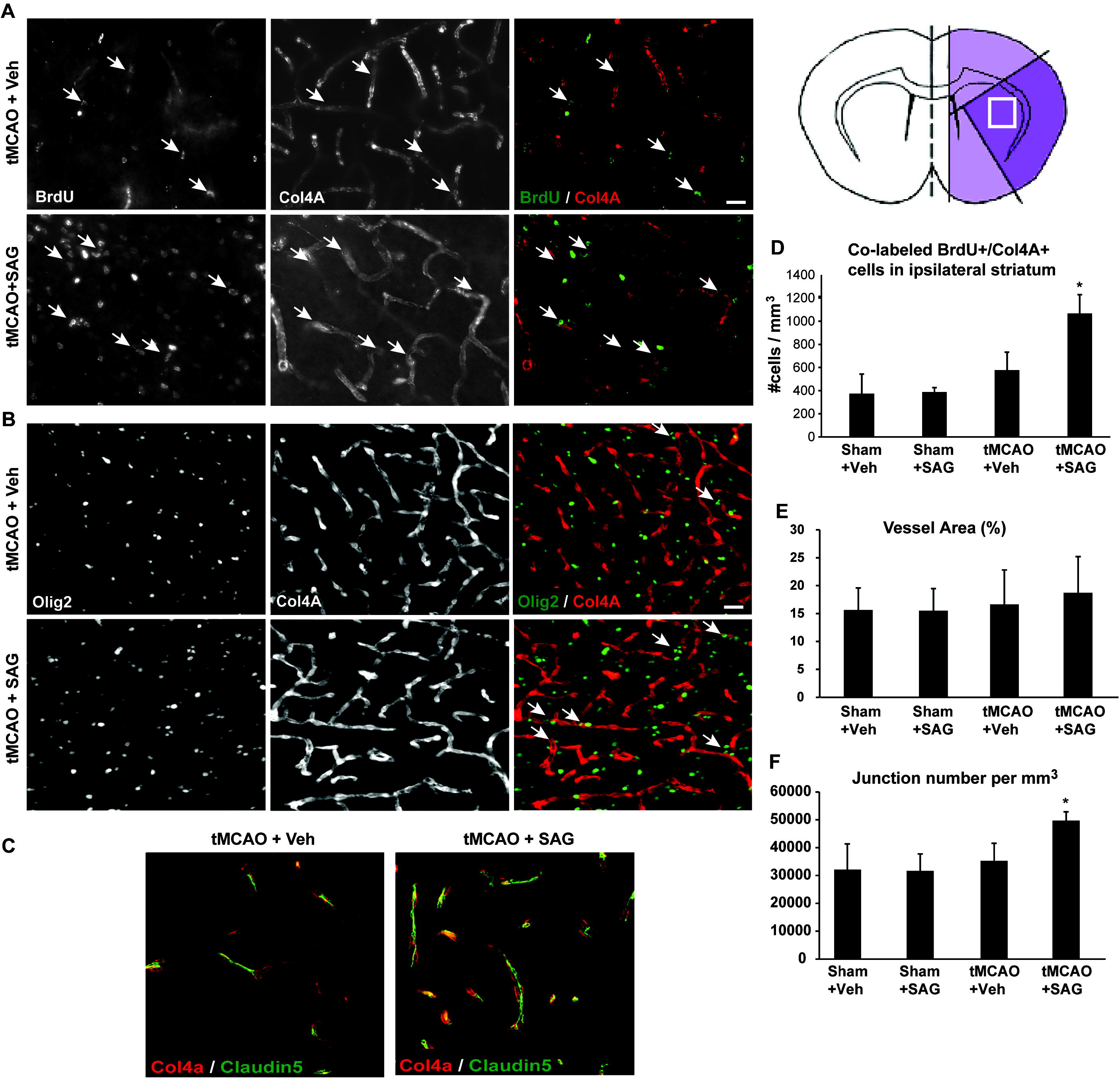
SAG enhances angiogenesis following tMCAO. (A) Cells in the injured core (striatum) stained with BrdU and Collagen 4A (Col4A), an EC marker, to visualize proliferation of cells associated with the vasculature. Arrows denote BrdU+ cells in direct contact with Col4A+ vessels. Scale bar, 100μm. Right, diagram depicting area of analysis. (B) Cells co-stained with Olig2 and Col4A in the striatum. Arrows denote Olig2+ cells in contact with Col4A+ vessels. (C) Cells co-stained with Col4A and Claudin5, a marker of blood brain barrier (BBB) structure. (D) Quantification of BrdU+/Col4A+ cells in the ipsilateral striatum per mm^3^. (E) Quantification of vessel area from Col4A-labeled vasculature in the motor and somatosensory cortical regions bordering the striatal core. (F) Quantification of vessel junction number per mm^3^ in the same region as (E). *, p < 0.05, one-way ANOVA with Tukey’s multiple comparisons test.

We observed a significant increase in the density of co-labeled, newly generated ECs in tMCAO+SAG animals (1067 ± 163 cells per hpf) versus tMCAO+Veh animals (578 ± 154, p<0.05; Fig. 5D), although there was no difference in vascular coverage (Fig. 5E), which could be attributed to the relatively early timepoint chosen for analysis(25). To determine whether endothelial proliferation contributed to increased angiogenesis, we analyzed vascular network features across cortical, striatal and white matter regions, and found a significant increase in vessel branching in tMCAO+SAG animals, consistent with endothelial proliferation at this early time point (tMCAO+SAG = 49447 ± 3189, tMCAO+Veh = 35068 ± 6268; p<0.05; Fig. 5F). Given these findings, it is possible that SAG-induced OPC proliferation is associated with the pro-angiogenic response.

### SAG improves long-term histology and cognitive function after tMCAO

To determine the long-term efficacy of single-dose SAG treatment after neonatal forebrain ischemia-reperfusion injury, we repeated the P10 tMCAO experiment and analyzed this cohort in young adulthood (Fig. 6A). At 8 weeks following tMCAO (P66) we used Cylinder Rearing (CR) testing to assess sensorimotor function and Novel Object Recognition (NOR) as a test of cognitive function and recognition memory. Brains were then harvested for volume analysis (P72). As shown (Fig. 6B), while there was no significant improvement in sensorimotor function after SAG treatment, we observed a significant improvement in cognitive function and recognition memory as assessed by NOR (Fig. 6C). There was a significant decrease in the percentage of novel object interaction in tMCAO+Veh compared to tMCAO+SAG animals, which did not differ from shams (tMCAO+Veh=0.44±0.04, tMCAO+SAG=0.58±0.03, p<0.05). Finally, we observed a persistent improvement in hemispheric brain volume at this later time point with SAG treatment (tMCAO+SAG=0.56±0.14, tMCAO+Veh=0.73±0.12, p<0.05; Fig. 6D), along with improved CC volume (tMCAO+Veh = 0.52 ± 0.06, tMCAO+SAG = 0.79 ± 0.06, p<0.05; Fig. 6E, F).

**Figure 6:**
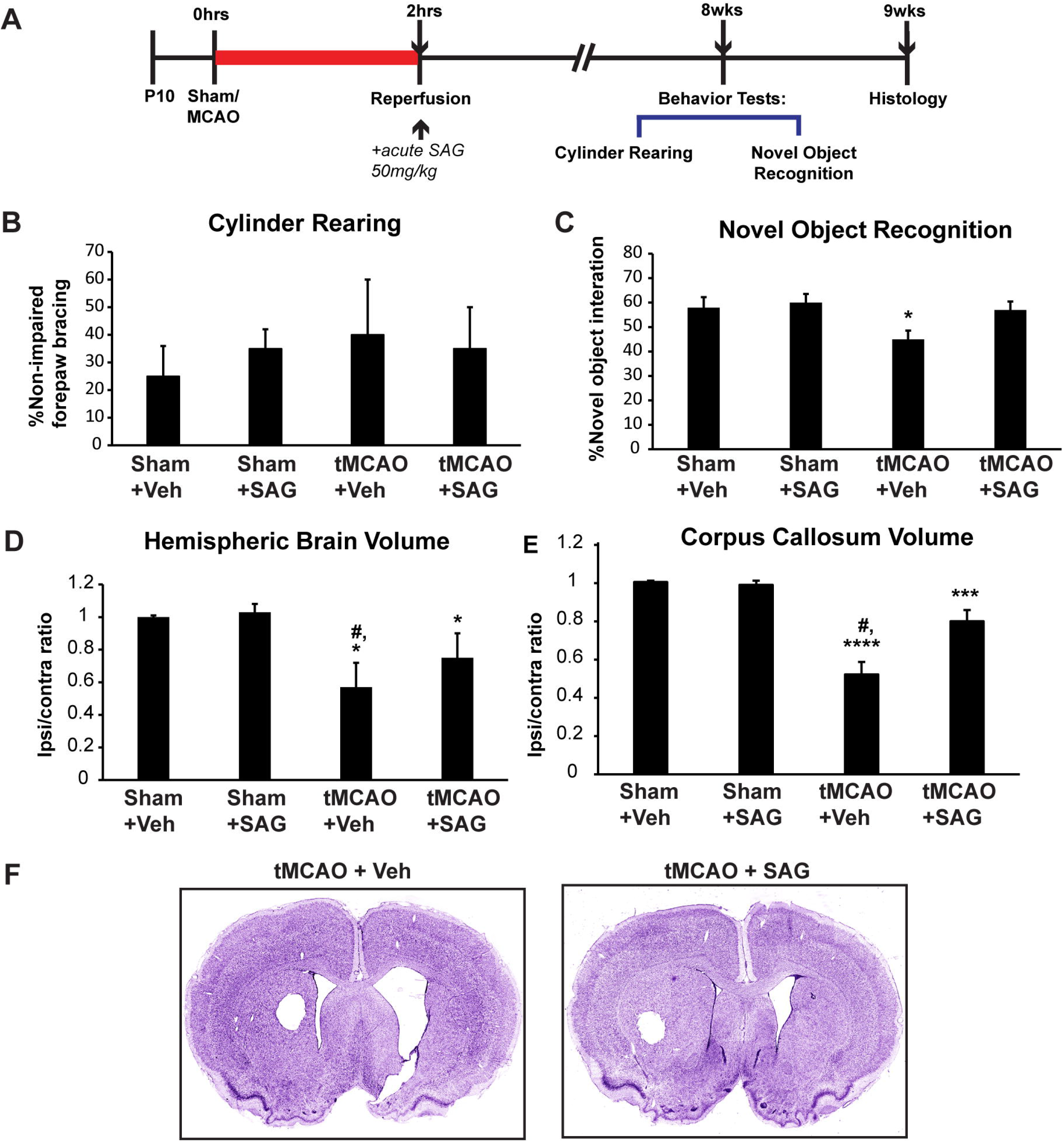
SAG shows improvements in behavioral tests. (A) Experimental timeline for surgery, treatment and behavioral tests for long-term neurobehavioral outcome. (B) Quantification of gross sensorimotor function between groups as assessed by forelimb preference in the Cylinder Rearing test. (C) Quantification of Novel Object Recognition test. (D) Quantification of ipsilateral-to-contralateral ratio of hemispheric brain volume. (E) Quantification of ipsilateral-to-contralateral ratio of corpus callosum volume. (F) Representative images of coronal sections from rat brains two months post-tMCAO or sham surgery. *, p<0.05, ***, p<0.001, ****, p<0.0001 vs Sham+Veh; #, p<0.05 vs tMCAO+SAG, one-way ANOVA with Tukey’s multiple comparisons test.

## DISCUSSION

In normal development, the Shh-signaling pathway is crucial for CNS patterning and cell fate specification(26), proliferation of cerebellar granule neuron precursors(27), neural circuit and synapse formation(28), vascular development(29) and oligodendrogenesis(30). We and others have shown benefits of Shh pathway activation in rodent models of neonatal cerebellar injury, Down syndrome (8) and adult stroke(13). Here, we extended these findings to a rat model of NS and report the novel finding that a single dose of SAG has beneficial effects to preserve brain volume, decrease inflammation, increased endothelial and oligodendroglial lineage proliferation and enhance long-term behavioral outcomes.

### SAG promotes oligodendroglial and endothelial proliferation after brain injury

We examined the effect of post-injury SAG administration on oligodendroglial lineage in two well established preclinical brain injury rodent models, tMCAO neonatal stroke to examine effects in developing brain injury and ethidium bromide (EB)-induced focal demyelination in adult rats to focus on the oligodendroglial response(12). There was a proliferative oligodendroglial response with SAG treatment in both models. Recent work has shown that increased proliferation is a prerequisite for the oligodendroglial differentiation(31) and that newly generated oligodendrocytes exhibit enhanced and accurate axonal myelination(32). Although we did not see a significant increase in mature CC1+ cells at these early time points, further studies are needed to determine whether SAG-induced oligodendroglial proliferation might benefit myelin generation in the long term.

Angiogenesis is critical for long-term repair following early focal ischemia-reperfusion injury, but remarkably little is known regarding the local vascular response to early focal ischemia-reperfusion injury in the brain(33). In an adult stroke model, Shh administration increased microvessel density and EC number 7-days after MCAO(13), while Smo agonist cyclopamine blocked neuroprotective effects in early subarachnoid hemorrhage(34). In our model of neonatal tMCAO, SAG administration promoted OPC numbers and angiogenesis. We found newly generated ECs in the injured core coupled with increased vascular branching after SAG treatment, suggesting enhanced vascular remodeling. Although we note that SAG did not regulate endothelial tight junction based on Claudin 5 expression, we do not rule out SAG affecting blood brain barrier permeability. Future extravasation studies at different time points will provide insight into this.

In the developing embryonic and postnatal brain, the vasculature engages in close interactions with migratory OPCs in both mice and humans(24) likely involving Wnt signaling during development and the response to injury(4, 24). In NS, we observed increased numbers of oligodendroglial cells in close apposition to blood vessels near the injured core in SAG-treated animals. Whether SAG directly promotes vascular remodeling or is a secondary consequence of oligodendrogenesis remains to be elucidated.

### One-time SAG treatment preserved brain volume and improved behavioral outcomes long-term after neonatal brain injury

We found that a single dose of SAG improved long-term behavioral outcomes and brain volume in a model of NS. Novel Object Recognition (NOR) testing showed treated animals had improved recognition memory of objects over a length of time. In contrast, we did not see long-term changes in sensorimotor function at young adulthood by CR, however, there was significant variability within groups, and this may test may not adequately analyze fine motor deficits at this time point following early injury. In addition, SAG treatment significantly preserved brain volume indicating that it has both short- and long-term benefit in a model of NS.

### Potential therapeutic benefits of SAG in NS

NS constitutes a major health problem in the United States and worldwide, with the majority of survivors exhibiting long-term neurodevelopmental deficits(35, 36). There are currently no accepted small molecule neuroprotective interventions for NS, and such therapies could be useful in acute settings outside major centers that provide hypothermia, including low- and middle-income countries.

Investigation of new treatment strategies for NS should take into account both etiology, and timing of injury to be as relevant as possible to human newborn injury and thus translation to the clinic(37). While erythropoietin has demonstrated consistent benefit with post-injury treatment in different models of early brain injury, including stroke(38), we note that multiple doses must be administered and that single dose therapy does not confer long-term benefits(39). In contrast, our findings demonstrate clear benefits of one-time treatment with SAG following tMCAO. However, as we have measured SAG bioactivity duration in a transgenic mouse model and not directly in our rat model, we acknowledge species differences may result in differential responses. In addition, we note that prolonged bioactivity does not imply that a single dose is either sufficient or most beneficial. Future work will examine alternative treatment plans such as multiple doses or delayed therapy.

Mounting evidence suggests that SAG is neuroprotective against injuries affecting preterm and term human neonates. We found that SAG preserves brain volume, increases OPC number, vascular proliferation, and long-term histological and cognitive benefit with a single dose given immediately post-NS. Therefore, this study supports investigation of SAG efficacy in larger animal models and further translational development with a view to ultimate testing in human clinical trials.

## Acknowledgements

We would like to thank members of the Gonzalez, Franklin and Rowitch labs for suggestions and technical assistance with experimental setup.

## Author contributions

VN, MC, AL, SK, and GG performed experimental procedures. VN, DHR and FG designed research and planned all the experiments. VN, MC, SK, GG, and FG analyzed the data and prepared the figures. VN, RJMF, DHR and FG wrote the article. DHR and FG conceived and led the project. All the authors read and approved the final version of the paper.

