## Supplementary Figure for "The neuroprotective effects of Sonic hedgehog pathway agonist SAG in a rat model of neonatal stroke"

**Supplemental Material**

**Supplementary Figure S1: Animal usage flow chart**

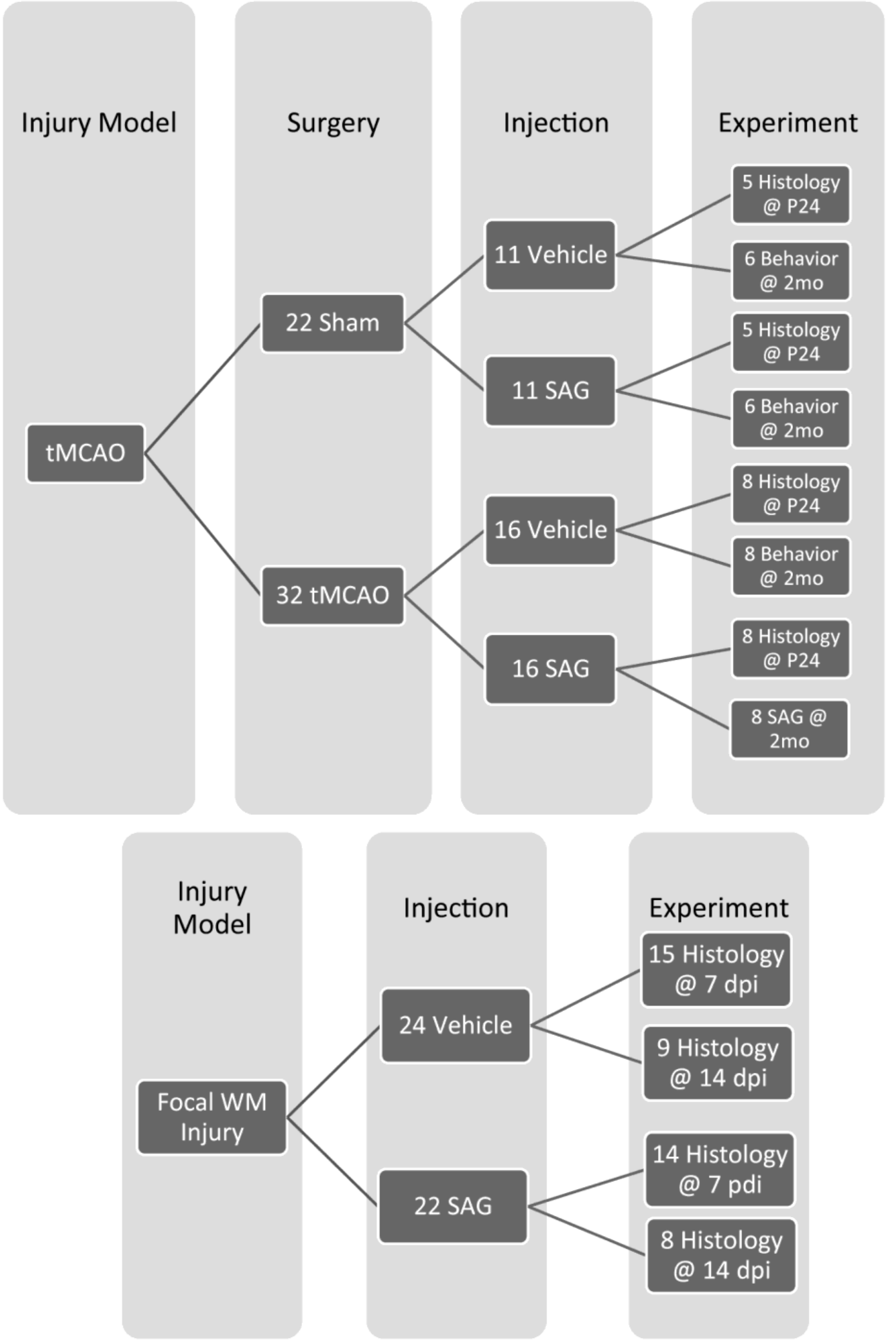


Total number of animals used for tMCAO and focal demyelination experiments, and their allocations.

**Supplementary Figure S2: Duration of SAG bioactivity *in vivo***

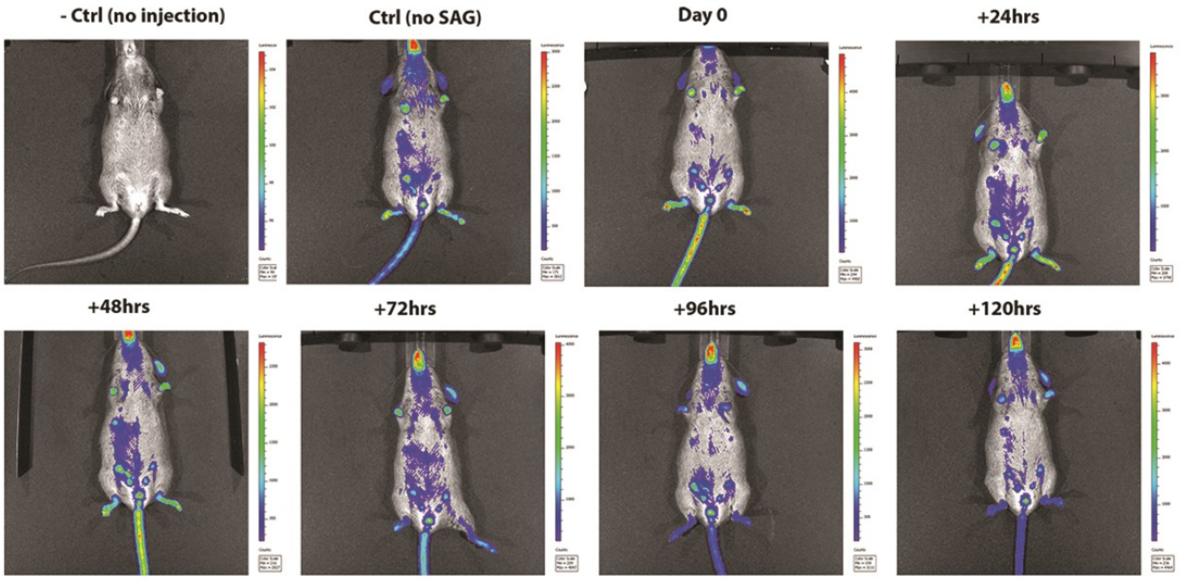


SAG injection (50mg/kg body weight) into *Gli-Luciferase* reporter mice shows increase in Shh downstream target *Gli1* in the limbs and tail lasting between 48 and 72 hrs.
